# Alcohol-induced sleep dysregulation in Drosophila is dependent on the neuropeptide PDF

**DOI:** 10.1101/2024.05.05.592572

**Authors:** Maria E. Ramirez-Roman, Nicolás L. Fuenzalida-Uribe, Genesis Ayala-Santiago, Jose L. Agosto, Alfredo Ghezzi

**Author notes:** **Correspondence:** Alfredo Ghezzi.

## Abstract

Alcohol exposure is known to trigger homeostatic adaptations in the brain that lead to the development of tolerance and dependence. These adaptations are also believed to be the root of a series of disturbances in sleep patterns that often manifest during the development of alcoholism and can have significant clinical and economic consequences. Unfortunately, the neuronal and genetic pathways that control the effects of alcohol on sleep are currently unknown, thus limiting our efforts to find effective treatment. In this study, we conduct a mechanistic exploration of the relationships between alcohol and sleep alterations using a *Drosophila* model system. We show that the genetic manipulation of the ventral lateral neurons (LNv) —a set of neurons known to control sleep in *Drosophila*— disrupts alcohol sensitivity and tolerance. Moreover, we show that alcohol exposure induces a series of alterations in sleep patterns that last for several days. Our results demonstrate that a single alcohol exposure promotes daytime sleep, alters the structure of sleep during the night, and reduces morning anticipatory behavior. In addition, we show that some of these alterations partially depend on the activity of the neuropeptide PDF, a key element in regulating sleep architecture. We propose that alcohol-induced sleep disruption stems from alterations in the activity of the PDF-releasing LNv neurons and that these alterations are similar to those that produce alcohol tolerance.

## Introduction

Alcohol use disorder is a severe condition and public health concern that influences the lives of patients, their families, and their communities. The 2017 National Survey on Drug Abuse and Health (NSDA) reports that 19.7 million Americans struggle with an issue associated with drug abuse, out of which 74% suffer from alcohol use disorder (SAMHSA, 2017). Accumulating evidence in both humans and in animal models suggests that alcohol use also contributes to a wide range of sleep disturbances, many of which have severe repercussions on health (Thakkar *et al*., 2015). In recovering alcoholics, this is especially problematic as these changes can last for prolonged periods, even after years of abstinence (Brower & Perron, 2010). Moreover, these sustained effects on homeostatic sleep processes can lead to relapse, as many individuals mistakenly turn to alcohol to induce sleep (Brower & Perron, 2010).

In humans, alcohol affects both the quantity and quality of sleep. However, these effects may differ depending on the dose, the mode of consumption, or even the sleep stage. For example, at acute high doses, alcohol is known to have sedative properties and can initially reduce sleep latency or the time it takes to fall asleep. However, as sleep progresses and the alcohol is metabolized, the organism may experience a state of episodic wakefulness and fragmented sleep, resulting in an overall decrease in sleep (Roehrs & Roth, 1995). While we know a substantial amount about the effects of alcohol on sleep patterns, little is known about the molecular mechanisms underlying the neuroadaptations that link both.

In recent years, the fruit fly, *Drosophila melanogaster*, has been used as a biological model to understand the neuronal and molecular basis of both sleep and alcohol use disorders. Extensive analyses have shown that sleep in fruit flies shows most of the fundamental features that characterize sleep in mammals, including sustained periods of inactivity, increased arousal threshold, and a homeostatic rebound period experienced upon sleep deprivation (Hendricks *et al*., 2000; Shaw *et al*., 2000). Similarly, fruit flies display alcohol responses that closely resemble those in humans. Flies develop tolerance and dependence to alcohol and can even display withdrawal symptoms after alcohol is withheld (Cowmeadow *et al*., 2006; Ghezzi *et al*., 2014; Rothenfluh *et al*., 2014; Scholz *et al*., 2000). Due to its powerful genetics and the myriad transgenic tools designed to explore distinct neural circuits, studies in this model system have resulted in significant advances in our understanding of the molecular underpinnings of these behaviors. In both cases, several distinct neural circuits and genetic pathways have now been identified, and many possible genetic interactions that link sleep and alcohol responses have started to emerge.

In both invertebrates and mammals, sleep is a tightly regulated process. It is well established that sleep timing is determined by the circadian clock, which controls the unfolding of rest/activity periods. However, sleep intensity and duration are believed to be controlled by homeostatic processes, where the duration of wakefulness influences the drive to sleep. While there is a close link between the circadian and homeostatic processes, these are believed to be controlled by separate mechanisms (Allada et al., 2017; Daan et al., 1984). Drosophilae have approximately 150 neurons that express the core molecular clock components and serve to regulate circadian rhythms of behavioral activity (King & Sehgal, 2020). These neurons are distributed in discrete clusters throughout the fly brain and express different neurotransmitters, neuropeptides, and receptors, serving different functional roles (Reviewed in Dubowy and Sehgal (2017)). Amongst these, the Drosophila ventral lateral neurons form a set of four or five small cells and four large cells within the accessory medulla, and they innervate the optic lobe and the dorsal protocerebrum (Shafer & Yao, 2014). These cells are known for releasing the neuropeptide PDF and controlling distinct aspects of circadian behavior and sleep (Chung et al., 2009; Parisky et al., 2008; Shang et al., 2008; Sheeba et al., 2008b). The neuropeptide PDF serves as a signaling molecule involved in synchronizing clock cells with each other, orchestrating behavioral activity, and regulating sleep. This peptide is expressed in all Drosophila ventral lateral neurons except for a single PDF-negative cell called “the 5th small LNv” (Helfrich-Förster, 1995).

Like sleep, alcohol neuroadaptations, such as tolerance, dependence, and withdrawal, are believed to be rooted, in part, in homeostatic adaptations elicited to counteract the sedative effects of alcohol (Ghezzi & Atkinson, 2011; Koob & Le Moal, 1997; Littleton, 1998). Studies of the homeostatic process behind alcohol use disorders in *Drosophila* have uncovered a series of mechanisms involved in regulating neural activity and synaptic transmission. Interestingly, many of the genes identified so far are also well-known sleep or circadian core and output genes (Park *et al*., 2017; Pohl *et al*., 2013). One example is the gene *slo*, which encodes a BK-type calcium-activated potassium channel (Atkinson *et al*., 1991). In both flies and humans, this ion channel protein is known to regulate neuronal activity and has been implicated in the development of tolerance to alcohol (Cowmeadow *et al*., 2006; Ghezzi *et al*., 2014; Scholz *et al*., 2000). Moreover, BK channels also appear to act as an output of the central circadian pacemaker cells. In flies, a loss-of-function mutation in *slo* produces circadian arrhythmia (Ceriani *et al*., 2002) and a loss of PDF neuropeptide signal (Fernandez *et al*., 2007).

Interestingly, a previous study exploring the neural substrates of alcohol tolerance in *Drosophila* has identified the PDF-releasing ventral lateral neurons (LNv) as critical elements in regulating alcohol sensitivity (Ghezzi *et al*., 2013). This study showed that targeted over-expression of the *slo* gene in LNv neurons significantly increases alcohol sensitivity, directly linking a known molecular mechanism associated with alcohol responses with a cellular pathway involved in sleep regulation. Thus, in this study, we decided to characterize the effects of alcohol on sleep in *Drosophila* and explore the role of the PDF-releasing LNv neurons in the relationship between alcohol and sleep.

## Methods

### Fly maintenance and stocks

All *Drosophila* stocks used in this study were raised on standard cornmeal agar medium (Nutri-Fly® Bloomington Formulation, Genesee Scientific) and kept in an incubator set at 25 °C, 80% relative humidity, and under a 12-hour light / 12-hour dark cycle. For all assays, newly enclosed flies were collected over a 2-day interval and studied 3–5 days after collection. Stocks used in this study were obtained from the Bloomington *Drosophila* Stock Center at Indiana University (Bloomington, IN) unless specified otherwise. The following lines were used for experiments: the wild-type line CS; the *Pdf ^01^* mutant line, which was backcrossed into the CS background for six generations to reduce genetic background effects as described in Pohl et al. (2013) (kindly provided by Dr. Nigel Atkinson from the University of Texas at Austin); the Pdf-GAL4 line (stock # 80939); the UAS-Kir2.1 line (stock # 6595); the UAS-TeTxLC line (stock # 28837); and the UAS-NaChBac line (stock # 9469). For Gal4 induction of UAS transgenes, female virgin Pdf-GAL4 flies were crossed with UAS-Kir2.1, UAS-TeTxLC & UAS-NaChBac male flies to inactivate/activate LNv neurons. Parental controls for transgenic experiments were generated by crossing either the Pdf-GAL4 driver line or the UAS-responder lines with the wild-type strain CS.

### Alcohol exposure

Alcohol exposure was performed using a custom-built alcohol delivery apparatus. The apparatus consisted of an aquarium air pump attached to two independent 30 ml midget bubblers (Ace Glass Inc.; Vineland, NJ) containing 10 ml of water or 95% alcohol. Airflow regulators were connected to each bubbler so that the flow of air passing through each bubbler could be controlled independently. Airflow from each bubbler was directed to perforated 15 ml or 50 ml conical tubes (depending on the number of flies being treated). For alcohol exposure, groups of flies were placed in the perforated conical tubes and exposed to a stream of humidified air at 1.5 LPM for 5 minutes to acclimate to the chamber and the airflow. After the acclimation period, the airstream was switched to alcohol-saturated air (1.5 LPM) for 20 minutes until all flies lost postural control. After the sedation period, the airstream was switched back to humidified air for 5 minutes to clear the alcohol from the chamber, and flies were subsequently returned to food for a 24-hour recovery period. All alcohol exposures were performed at ZT 9 or approximately 3 hours before lights off.

### Analysis of alcohol sensitivity and tolerance

Alcohol sensitivity and tolerance experiments were performed over the course of two days. On the first day, age-matched female flies were divided into two groups —one (the alcohol group) was exposed to alcohol-saturated air, and the other (the control group) was exposed to humidified air. For alcohol treatment, approximately eight flies from the alcohol group were placed in a perforated 15 ml conical tube and exposed to alcohol as described above. For the mock-exposure control groups, flies were simultaneously exposed to humidified air for the entire 30 minutes (1.5 LPM) using an identical setup with no alcohol. On the second day, individual flies from both groups were placed in *Drosophila* Activity monitors (DAM2) equipped with a gas manifold (MAN2), which allows tracking of the activity of individual flies during exposure to the alcohol-saturated air stream (Trikinetics; Waltham, MA). In this case, flies from both groups were exposed to the alcohol-saturated air stream (1.5 LPM) until all flies were sedated. Activity bouts were automatically recorded by the DAM2 every minute during the course of alcohol exposure. The time of sedation for each fly was calculated by identifying the time of the last movement detected by the monitor. For statistical analysis, the average sedation time was calculated for each group. Outliers whose average deviated more than two standard deviations from the overall averages were removed. Sensitivity to alcohol across genotypes was determined by comparing the time of sedation of flies in the control groups. Alcohol tolerance was calculated by comparing the time of sedation between the control group and the alcohol group.

### Sleep Analysis

Prior to sleep analysis, age-matched female flies were divided into two groups —one (the alcohol group) was exposed to alcohol-saturated air, and the other (the control group) was exposed to humidified air. For alcohol exposure, approximately 32 flies from the alcohol group were placed in a perforated 15 ml conical tube and exposed to alcohol as described above. For the mock-exposure control groups, flies were simultaneously exposed to humidified air for the entire 30 minutes (1.5 LPM) using an identical setup with no alcohol. After treatment, flies were transferred to individual 65 mm x 5 mm glass tubes (Trikinetics, Waltham, MA) containing fly food. Tubes were then placed in *Drosophila* Activity Monitors (DAM2) within an environmentally controlled incubator set at 25 °C, 80% relative humidity, and under a 12-hour light / 12-hour dark cycle. Locomotor activity was monitored using the Trikinetics monitoring system (Waltham, MA) for a period of 5 to 9 days. Locomotor activity was collected in 1 min bins as previously described (Agosto *et al*., 2008). Sleep was measured as bouts of uninterrupted 5 min of inactivity. Daily sleep parameters were analyzed using MATLAB software (Natick, MA) as described in (Parisky *et al*., 2008). Total sleep duration, sleep latency, number of sleep episodes, mean sleep episode duration and total locomotor activity were analyzed for each 12-hour period of the light/dark protocol per day. Morning and evening anticipation indices and the night sleep ratios were calculated using the Rethomics framework in R (Geissmann *et al*., 2019) as described in Harrisingh et al. (2007). For morning anticipation, the sum of activity for each fly between ZT21 and ZT24 was divided by the sum of activities between ZT18-24 for each day analyzed. For evening anticipation, the sum of activity for each fly between ZT9 and ZT12 was divided by the sum of activities between ZT6-12 for each day analyzed. For night sleep ratios, the sum of total sleep for each fly between ZT18 and ZT24 was divided by the sum of activities between ZT12-18 for each day analyzed.

### Statistical Analysis

For all assays performed, statistically significant differences were calculated using GraphPad Prism software. For alcohol tolerance, significant differences in sedation time between the first and second alcohol exposures were determined using a standardized Student’s t-test. This test was conducted separately for each genotype studied, as these are fully independent assays. To determine differences in alcohol sensitivity between genotypes, statistical comparisons of the sedation time during the first exposure were performed between each experimental line and the respective control lines. For *Pdf* ^01^, sedation time was compared to that of CS, and significant differences were determined using a standardized Student’s t-test. For, Pdf-GAL4;UAS-Kir2.1, Pdf-GAL4;UAS-TeTxLC, and Pdf-GAL4;UAS-NaChBac, sedation time was compared to the sedation time of the two corresponding parental control lines. In this case, significant differences were determined using an ordinary one-way ANOVA, with Bonferroni post-hoc test for multiple comparisons. For the analysis of the effects of alcohol on sleep parameters over time, the daily values of each measurement obtained were compared. Statistically significant differences between alcohol-treated and mock-treated flies were calculated for each genotype using two-way Repeated Measures ANOVA to obtain the main effect of the alcohol treatment, the main effect of time, and the interaction between treatment and time. A Bonferroni posthoc test for multiple comparisons was performed to calculate the significance of the treatment effect per day. Additionally, a three-way ANOVA was performed to estimate the impact of genotype on the differences between alcohol-treated and mock-treated flies. This allows a direct statistical comparison of all three variables: genotype, treatment, and time. A P value threshold of 0.05 was used for all tests to determine statistical significance. The P values, the number of subjects used for each test (n), and the name of each post-hoc test performed are detailed in the caption for the corresponding Figures.

## Results

### Alcohol sensitivity and tolerance are dependent on PDF

We began our experiments by analyzing the role of the PDF-releasing LNv neurons on alcohol sensitivity and tolerance. Alcohol sensitivity is defined as the baseline level of response that an organism displays to a particular effect of alcohol. In contrast, alcohol tolerance refers to the change in the sensitivity to alcohol after prior alcohol exposure. To measure these two endophenotypes of alcohol, we utilized a two-day alcohol exposure approach. On the first day, each genotype was divided into two groups: an experimental group that would be exposed to alcohol vapor and a control group that would be exposed to humidified air. On day two, we administered alcohol to both groups simultaneously and monitored the time to sedation for each group independently using an automated activity monitor. In essence, the alcohol administered on day two would be the first alcohol exposure for the control group and the second exposure for the experimental group. Differences in sensitivity between genotypes were, thus, quantified by comparing the time-to-sedation of flies after their first exposure to alcohol. In contrast, tolerance within each genotype was quantified by comparing the time-to-sedation between the first and second exposure to alcohol.

The first comparison we performed was between the wild-type CS flies and the *Pdf ^01^* line —a null mutant in the gene that encodes PDF neuropeptide (Figure 1A). The PDF neuropeptide is one of the outputs from the LNv neurons and a critical mediator of sleep regulation in flies (Park & Hall, 1998). This mutant has been previously backcrossed to the CS background for six generations to minimize genetic background effects (Pohl *et al*., 2013). We observed that CS flies took approximately 20 minutes to sedate after their first exposure, while the *Pdf ^01^* flies took nearly 40 minutes. The student’s t-test analysis reveals this difference was significant, suggesting a clear effect of the mutation on alcohol sensitivity. Moreover, when comparing the first and second exposure for each genotype, we observed that while CS flies acquired tolerance —they almost doubled the time-to-sedation in their second exposure— while the *Pdf ^01^*mutant did not acquire tolerance. In the latter case, the time-to-sedation between first and second exposures was not significantly different. This evidence suggests that the PDF neuropeptide is critical for both sensitivity and tolerance to alcohol.

**Figure 1.**
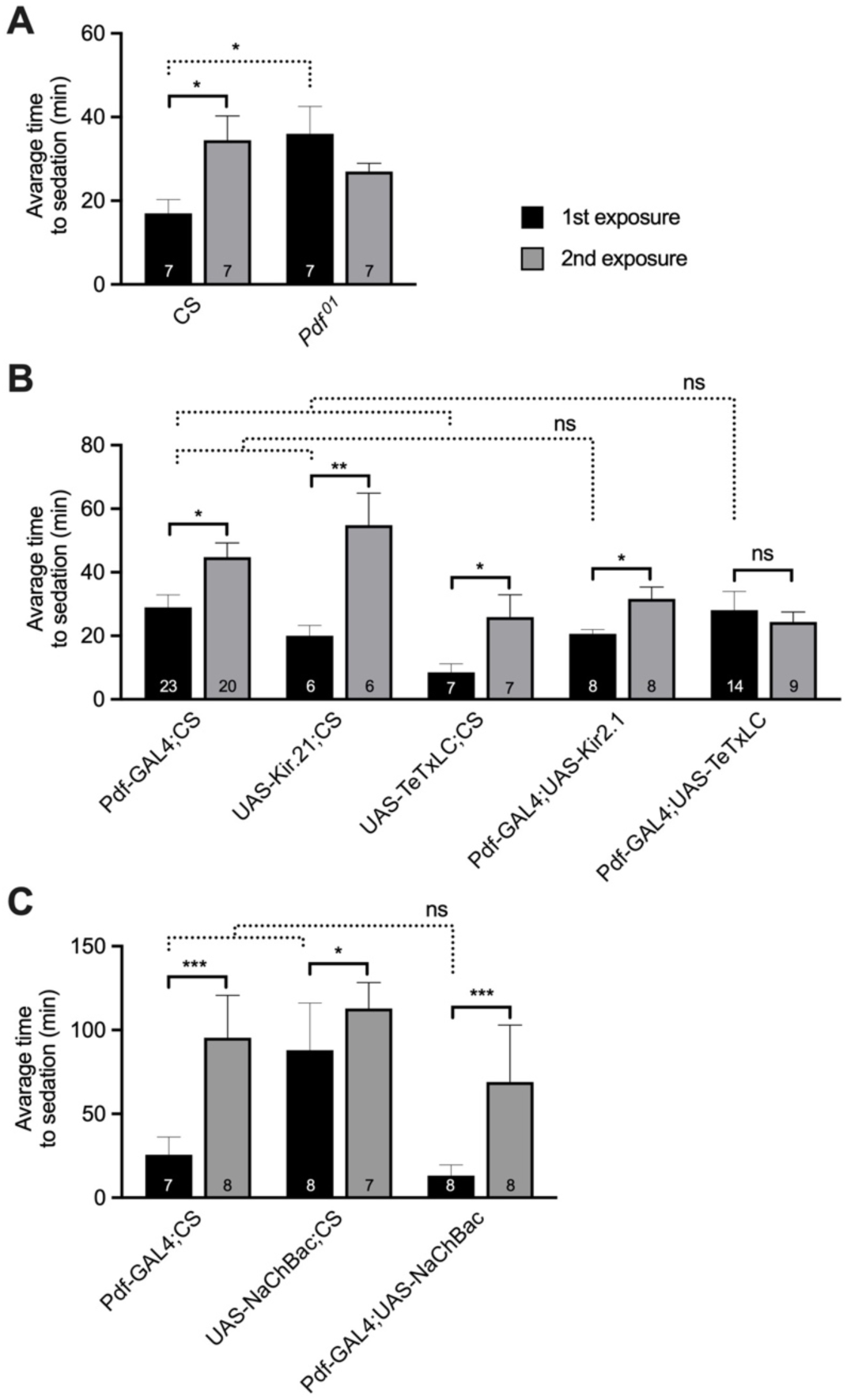
Alcohol sensitivity and tolerance depend on the pigment dispersing factor (PDF). A) The *Pdf*^01^ null mutation increases the average time to sedation after the 1st exposure to alcohol and blocks the capacity to acquire alcohol tolerance. The average time to sedation upon first or second sedation to alcohol is shown for wild-type Canton-S (CS) control flies and the *Pdf* ^01^ mutants. B) Inhibition of PDF-expressing neurons by expression of the tetanus toxin light chain (TeTxLC) protease or overexpression of a heterologous inward rectifying potassium channel (Kir2.1) differentially affects the capacity to acquire alcohol tolerance. The average time to sedation upon first or second sedation to alcohol is shown for parental control flies (Pdf-GAL4/CS, UAS-kir2.1/CS, UAS-TeTxLC/CS) and the genetically inhibited lines (Pdf-GAL4/UAS-Kir2.1 and Pdf-GAL4/UAS-TeTxLC). C) Activation of PDF-expressing neurons by expressing the heterologous sodium channel construct NaChBac augments the capacity to acquire alcohol tolerance. The average time to sedation upon first or second sedation to alcohol is shown for parental control flies (Pdf-GAL4/CS, UAS-NaChBac/CS) and the genetically activated lines (Pdf-GAL4/UAS-NaChBac). Error bars represent SEM. n for each genotype is displayed as a number within each bar. For alcohol tolerance, statistically significant differences between 1^st^ and 2^nd^ exposures were determined by Student’s t-test (continuous horizontal brackets): * denotes P < 0.05; ** denotes P < 0.01; *** denotes P < 0.001; ns = not significant. For alcohol sensitivity, statistically significant differences between genotypes were determined by Student’s t-test (in A) or one-way ANOVA (in B and C) (dotted horizontal brackets): * denotes P < 0.05; ns = not significant.

To better understand the role of the PDF-expressing neurons on alcohol sensitivity and tolerance, we used the UAS-GAL4 system to genetically alter neuronal activity in the LNv neurons (Brand *et al*., 1994; Hodge, 2009). Using a transgenic construct carrying the coding sequence of the yeast transcription factor GAL4 under the control of the *Pdf* tissue-specific enhancer, we were able to express two distinct ectopic proteins known to silence neural activity and one known to enhance neural activity. These proteins were tethered to the Gal4-responsive UAS sequence so that they could be specifically expressed in the LNv neurons using the Pdf-GAL4 driver. The first silencing approach is based on the electrical suppression of neural excitability by overexpression of the Kir2.1 inward rectifier potassium channel. This manipulation is highly effective at shunting the electrical activity of excitable cells in both mammals and *Drosophila* (Baines *et al*., 2001; Johns *et al*., 1999; White *et al*., 2001). The second approach is based on the expression of the tetanus toxin light chain (TeTxLC) protease. The TeTxLC protease cleaves synaptobrevin, syntaxin, or SNAP-25, thus blocking synaptic transmission and neuropeptide release (Ding *et al*., 2019; Sweeney *et al*., 1995).

Finally, we also tested a method for selectively enhancing neuronal excitability through overexpression of the voltage-activated bacterial sodium channel NaChBac (Ren *et al*., 2001). This method has been shown to enhance excitability in *Drosophila* neurons, including the LNvs (Luan *et al*., 2006; Sheeba *et al*., 2008a).

When subjected to the alcohol sensitivity and tolerance tests, we observed distinct effects induced by the two silencing methods (Figure 1B). First, while there was no significant effect of silencing on baseline alcohol sensitivity when compared to the respective parental controls, it became evident that sensitivity was highly variable across all genotypes, suggesting that alcohol sensitivity is distinctly susceptible to genetic background. However, when looking at alcohol tolerance —by comparing time to sedation between first and second alcohol exposures within each genotype— it was immediately apparent that all genotypes were able to acquire tolerance except for the group where LNv synaptic transmission was blocked with TeTxLC. In contrast, activation of LNv neurons using NaChBac showed robust tolerance to alcohol but no significant effect on baseline sensitivity (Figure 1 C). Together, these data suggest that alcohol tolerance is dependent on neuropeptide release by LNv neurons rather than on the electrophysiological state of the cells.

### Alcohol promotes daytime sleep in wild-type flies

Given the role of LNv neurons in regulating alcohol tolerance, we next sought to determine the effects of alcohol on the sleep profile of flies and explore the role of PDF-expressing neurons in the observed alcohol-induced sleep alterations. For this, wild-type CS flies and *Pdf* ^01^ mutants were subjected to a single alcohol exposure, and their activity and sleep patterns were evaluated for nine consecutive days using the *Drosophila* Activity Monitors (DAM2). In flies, sleep is defined as periods of locomotor inactivity lasting at least 5 minutes. These periods are associated with an increased arousal threshold, as assessed using mild mechanical stimulation (Hendricks *et al*., 2000; Shaw *et al*., 2000). Figure 2A shows the percentage of time that wild-type CS flies spend sleeping during the 12-hour light and 12-hour dark periods of every day after exposure to alcohol or the respective untreated controls. In contrast, Figure 2B shows the percentage of time that *Pdf* ^01^ mutant flies spend sleeping during the same period after exposure. As observed, during the first two days, there was a clear increase in daytime sleep after alcohol exposure in both fly lines tested, albeit this effect diminished as the days progressed. At night, however, there was no apparent effect of alcohol on the CS flies as both alcohol-treated and untreated flies seemed to sleep on average in equal amounts. This contrasted with what is observed in the *Pdf* ^01^ mutants, where alcohol-treated flies slept more than untreated flies on the first night but less than untreated flies on the following nights.

**Figure 2.**
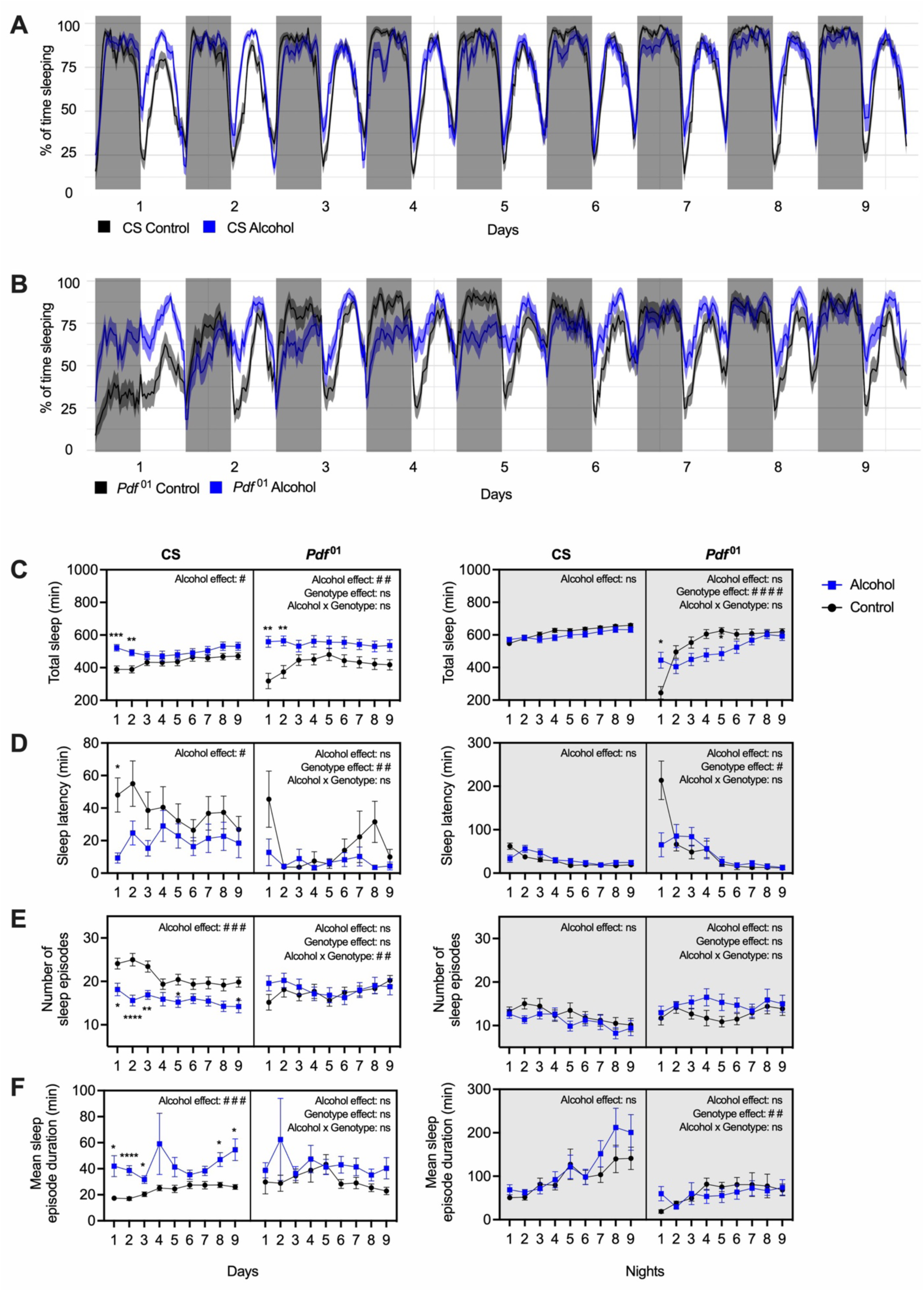
A single alcohol exposure differentially alters the activity pattern and sleep architecture of wild-type and *Pdf* ^01^ mutant flies. A) The percentage of time spent sleeping (in 30-minute bins) of control (black line) and alcohol-exposed (blue line) wild-type flies over 9 consecutive days after exposure. B) Percentage of time spent sleeping (in 30-minute bins) of control (black line) and alcohol-exposed (blue line) *Pdf* ^01^ flies over 9 consecutive days after exposure. C-F) Sleep parameters for each genotype (CS and *Pdf* ^01^) for every daytime period (left) and night-time period (right) after alcohol exposure. Sleep parameters shown are as follows: C) total time sleeping, D) sleep latency, E) number of sleep episodes, and F) mean sleep episode duration. Averages per day are displayed for alcohol-treated flies (blue) and untreated controls (black). Error bars are SEM; n = 31 (CS-control), 30 (CS-alcohol), 27 (*Pdf* ^01^-control), 22 (*Pdf* ^01^-alcohol). Statistically significant differences between alcohol and control groups across each curve were determined by two-way ANOVA with Bonferroni multiple comparisons post-hoc test to establish individual differences for each day: * denotes P < 0.05; * denotes P < 0.01; *** denotes P < 0.001; **** denotes P < 0.0001; ns = not significant. Significant interactions between the effect of alcohol and genotype were determined by three-way ANOVA: # denotes P < 0.05; # # denotes P < 0.01; # # # denotes P < 0.001; # # # # denotes P < 0.0001; ns = not significant. Periods shaded in gray denote night-time, and unshaded periods denote daytime.

To provide a deeper understanding of the effects of alcohol on sleep architecture, we quantified specific sleep parameters for every day after alcohol exposure. The sleep parameters analyzed were: (I) the total sleep, described as the total amount of sleep that flies experienced during the day or the night; (ii) the sleep latency or the time it takes for the fly to fall asleep after the lights-on or lights-off transition; (iii) the number of sleep episodes, which is defined as the number of intervals of sleep the flies experienced during a specific period; and (iv) the mean sleep episode duration, which is calculated by adding the amount of time the flies spent sleeping in each sleep episode divided by the number of sleep episodes. Measurements for each of these parameters are shown in Figure 2C-F for every day (left panel) and night (right panel) after alcohol exposure in CS wild-type flies and *Pdf* ^01^ mutants.

For CS flies, our quantitative analysis revealed that the total sleep in alcohol-exposed flies increased significantly during the light period of days 1 and 2. Nevertheless, no significant differences were observed in total sleep during days 3, 4, 5, 6, 7, 8, or 9. During the dark period, however, there were no significant differences in total sleep across all nine days after exposure (Figure 2C). Daytime sleep latency was shorter in alcohol-exposed flies than controls during day one but gradually returned to control levels during the rest of the days. No differences were observed in the sleep latency between air and alcohol-exposed flies at night (Figure 2D). During the day, alcohol-exposed flies experienced more consolidated sleep, reflected by a decrease in the number of sleep episodes and an increase in their duration compared to controls across several days. No differences were observed in the number of sleep episodes and the mean sleep episode duration at night (Figures 2E and 2F, respectively).

Quantitative analysis of the *Pdf* ^01^ sleep pattern revealed that alcohol-exposed *Pdf* ^01^ flies slept more during the light period than controls for the first two days but returned to control levels after the third day. This was similar to the effect in CS. Interestingly, during the dark period, alcohol-exposed *Pdf* ^01^ flies slept significantly more than controls for the first day but quickly switched to sleeping less than controls after the second day. This pattern continued for three days but normalized to control levels after that (Figure 2C). This is markedly different from what was observed in wild-type flies, as alcohol had no significant effect on total night-time sleep. Daytime sleep latency was significantly reduced in *Pdf* ^01^ flies compared to wild-types but was unaffected by alcohol treatment. At night, however, sleep latency was increased in *Pdf* ^01^ flies as compared to wild-types, especially on the first night after flies were placed in the monitors, but this effect was suppressed by alcohol treatment (Figure 2D). Finally, no differences were observed in the number of sleep episodes or their duration between alcohol-treated *Pdf* ^01^ flies and the untreated controls. This contrasts with wild-type flies that displayed a long-term increase in the number of episodes of shorter duration during daytime sleep after alcohol exposure (Figures 2E and 2F).

### Alcohol exposure alters night sleep architecture

While we did not observe an effect of alcohol on total night-time sleep, closer inspection of the night sleep pattern revealed a significant alteration in the structure of night sleep after alcohol exposure. In particular, we observed that while wild-type control flies slept consistently well throughout the night, alcohol-exposed flies slept significantly less during the first half of the night than during the second half. In fact, alcohol-exposed flies slept longer in the last six hours of the night than control flies, which started to wake up in anticipation of the morning, approximately 3 hours before the lights-on transition. This difference can be observed in the average daily sleep over the nine days after alcohol exposure (Figure 3A). This effect was less pronounced in *Pdf* ^01^ flies, as both control and alcohol-exposed flies seemed to have slept equally well throughout the night. (Figure 3B).

**Figure 3.**
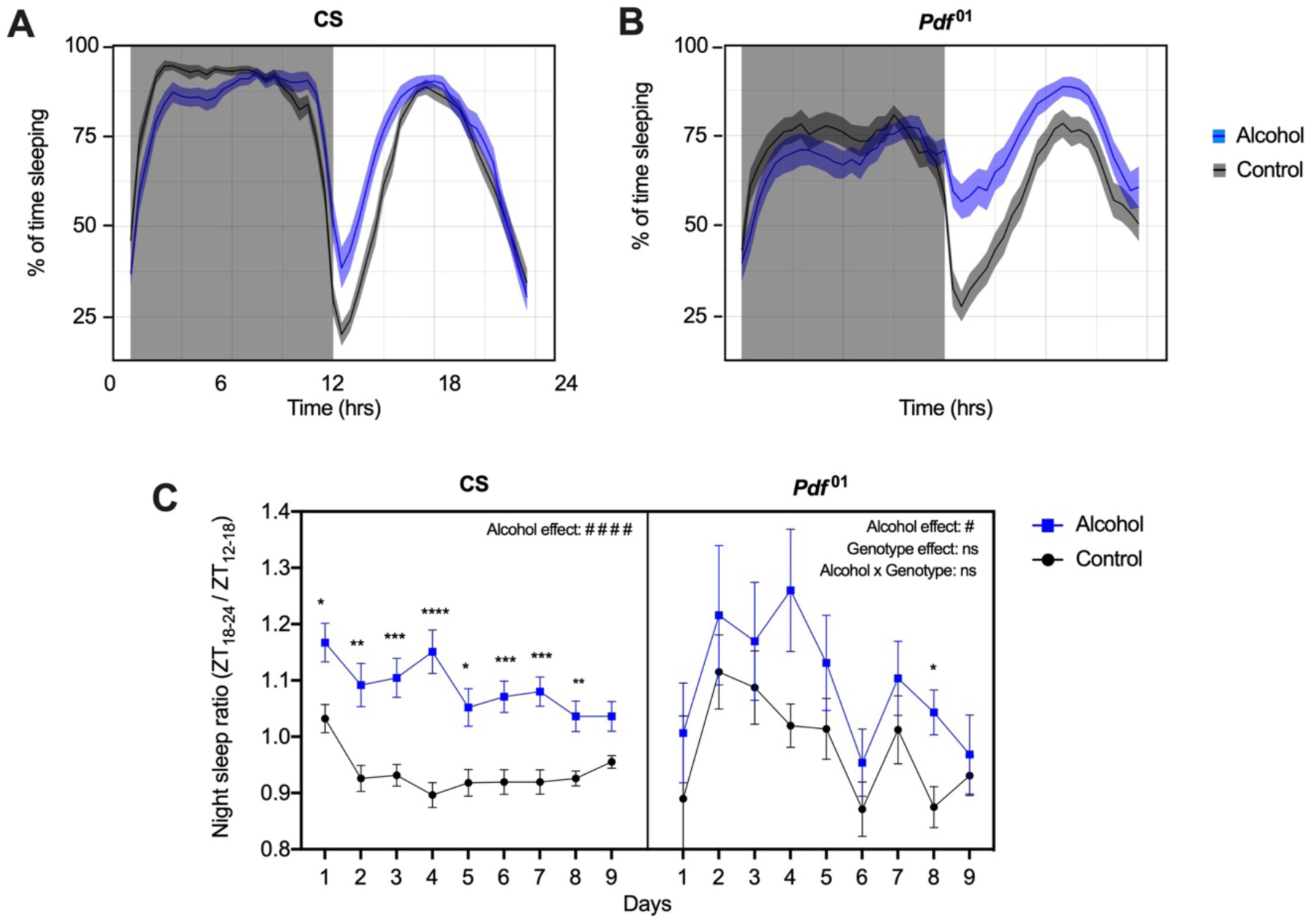
Alcohol exposure alters night sleep architecture. A) The average percentage of time sleeping (in 30-minute bins) of control (black line) and alcohol-exposed (blue line) of wild-type CS flies over a nine consecutive day period after alcohol exposure is displayed. B) The average percentage of time sleeping (in 30-minute bins) of control (black line) and alcohol-exposed (blue line) of *Pdf* ^01^ flies over a nine consecutive day period after alcohol exposure is displayed. C) The ratio of sleep displayed during the first 6 hours of the night over the sleep displayed in the last 6 hours of the night is shown for CS (left) and *Pdf* ^01^ mutant flies (right). Averages per day are displayed for alcohol-treated flies (blue) and untreated controls (black). Error bars are SEM; n = 31 (CS-control), 30 (CS-alcohol), 27 (*Pdf* ^01^-control), 22 (*Pdf* ^01^-alcohol). Statistically significant differences between alcohol and control groups across each curve were determined by two-way ANOVA with Bonferroni multiple comparisons post-hoc test to establish individual differences for each day: * denotes P < 0.05; * denotes P < 0.01; *** denotes P < 0.001; **** denotes P < 0.0001; ns = not significant. Significant interactions between the effect of alcohol and genotype were determined by three-way ANOVA: # denotes P < 0.05; # # denotes P < 0.01; # # # denotes P < 0.001; # # # # denotes P < 0.0001; ns = not significant. Periods shaded in gray denote night-time, and unshaded periods denote daytime.

To quantify this effect, we calculated the ratio of sleep displayed during the first 6 hours of the night over the sleep displayed in the last 6 hours. In wild-type CS flies, this ratio was consistently higher in alcohol-treated flies than in controls over the first eight days of the experiment. In contrast, *Pdf* ^01^ flies showed no significant differences between the two groups, indicating that this effect is partially dependent on the PDF neuropeptide (Figure 3C).

### Alcohol exposure disrupts morning anticipation

*Drosophila* display morning and evening locomotor activity peaks with robust anticipatory activity before the transitions of lights-on and lights-off (Dubowy & Sehgal, 2017). The observation that alcohol-treated flies slept beyond the point that control flies would typically start to wake during the last hours of the night prompted us to explore the effect of alcohol on the anticipatory behavior of light transitions. We collected and analyzed the locomotor activity patterns of flies over a period of nine days following alcohol exposure. Figure 4A shows the activity profiles during 12-hour light and 12-hour dark periods for every day after exposure to alcohol or the respective controls in a double-plotted actogram. For wild-type CS flies, the morning and evening anticipatory activity (red and gray arrows, respectively) became immediately apparent as a gradual increase in activity that started several hours before the light transition. This anticipation is evident for every single day plotted.

**Figure 4.**
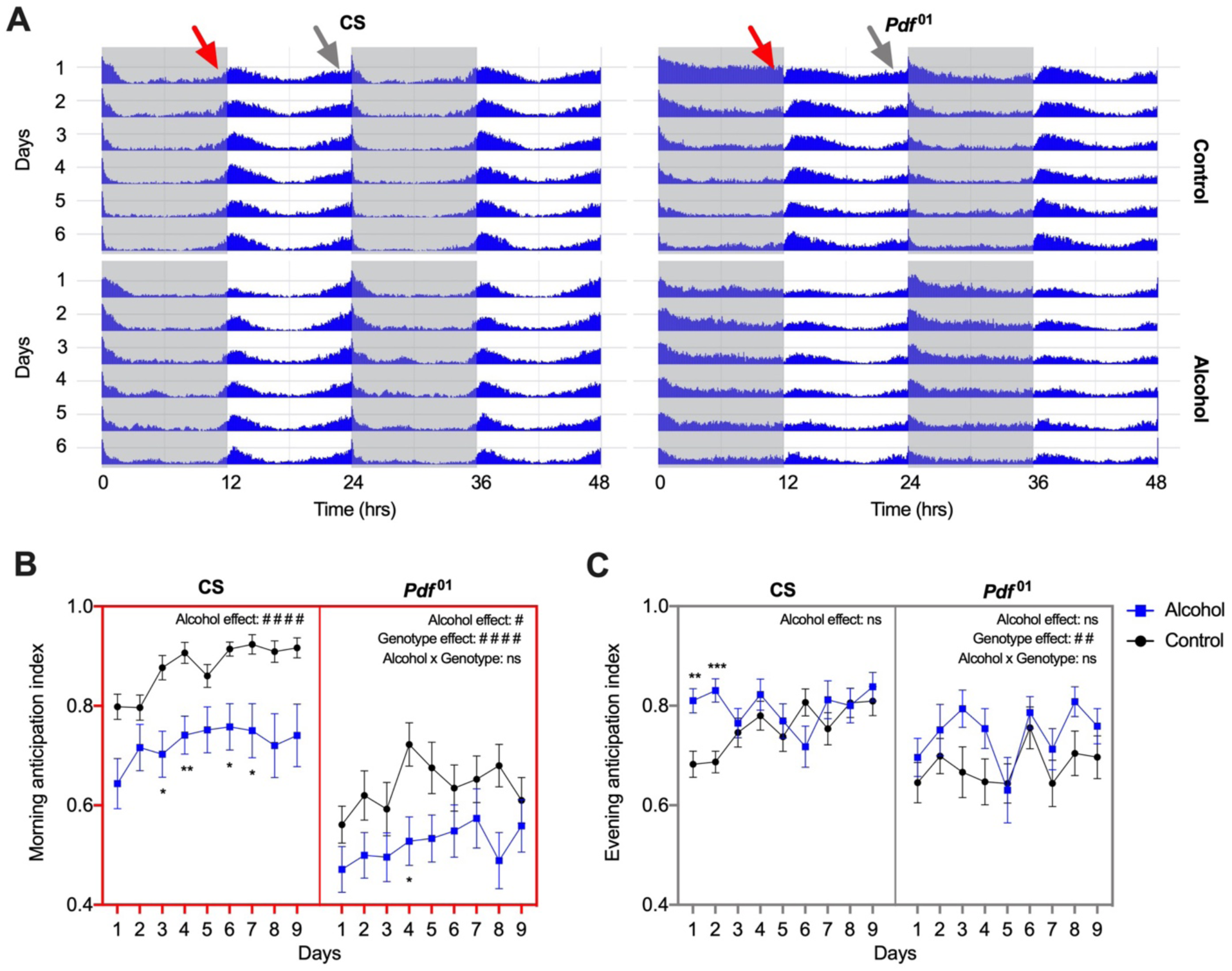
Alcohol exposure disrupts morning anticipation. A) Double-plotted actograms for wild-type CS (left) and *Pdf* ^01^ (right) of control (top) or alcohol-treated (bottom) flies are shown. Each row represents a different day after alcohol (or mock) exposure. Red arrows point at morning anticipatory activity. Gray arrows point at evening anticipatory activity. Periods shaded in gray denote night-time, and unshaded periods denote daytime. B) The morning anticipation index is shown for CS (left) and *Pdf* ^01^ mutant flies (right). Averages per day are displayed for alcohol-treated flies (blue) and untreated controls (black). C) The evening anticipation index is shown for CS (left) and *Pdf* ^01^ mutant flies (right). Averages per day are displayed for alcohol-treated flies (blue) and untreated controls (black). Error bars are SEM; n = 31 (CS-control), 30 (CS-alcohol), 27 (*Pdf* ^01^-control), 22 (*Pdf* ^01^-alcohol). Statistically significant differences between alcohol and control groups across each curve were determined by two-way ANOVA with Bonferroni multiple comparisons post-hoc test to establish individual differences for each day: * denotes P < 0.05; * denotes P < 0.01; *** denotes P < 0.001; ns = not significant. Significant interactions between the effect of alcohol and genotype were determined by three-way ANOVA: # denotes P < 0.05; # # denotes P < 0.01; # # # denotes P < 0.001; # # # # denotes P < 0.0001; ns = not significant. Periods shaded in gray denote night-time, and unshaded periods denote daytime.

We quantitatively analyzed the anticipatory behavior for both CS (Figure 4B) and *Pdf* ^01^ mutant flies (Figure 4C). Morning and evening anticipation indices were calculated for individual alcohol-treated and control flies using the activity ratio during the last three hours before the transition over the activity level of the six hours before the transition (Harrisingh *et al*., 2007). Our analysis revealed that in wild-type CS flies, morning anticipation was severely reduced after alcohol exposure and that this effect lasted for several days. In contrast, in the *Pdf* ^01^ mutant flies, morning anticipation was decreased in both alcohol-treated and untreated controls. For evening anticipation, alcohol induced a slight enhancement in CS flies that lasted for the first two days. However, there was no significant effect of alcohol in *Pdf* ^01^ mutant flies.

## Discussion

Alcohol is a well-known suppressor of neural activity, and as such, it can trigger homeostatic neuroadaptations to counteract the effects of the drug. These adaptations can lead to tolerance and dependence on alcohol and can even affect the normal function of several neuronal processes (Ghezzi & Atkinson, 2011; Koob & Le Moal, 2001; Littleton, 1998). Sleep is highly sensitive to internal and external factors and can be modulated accordingly to adapt to its environment or to satisfy the needs of the organism (Allada *et al*., 2017; Okamoto-Mizuno & Mizuno, 2012). It is, thus, not surprising that these two processes interact. Using *Drosophila* and a combination of mutants and genetic tools to manipulate the activity of neural circuits, we have shown that a small set of circadian clock cells known to regulate sleep and arousal states are also involved in the regulation of alcohol-induced responses in flies, further linking these two phenomena. Additionally, we have characterized, for the first time, the effects of acute alcohol exposure on sleep regulation in *Drosophila*.

In our initial analysis of alcohol responses, we showed that a mutation in the gene that encodes the PDF neuropeptide can significantly alter the baseline sensitivity to alcohol and eliminate the capacity of flies to develop tolerance to alcohol. In general, *Pdf* ^01^ mutants were more resistant to the effects of alcohol than wild-type CS flies. When looking at tolerance, however, we observed that *Pdf* ^01^ flies were not able to gain tolerance like the wild-type flies. This latter effect could result from the increase in baseline resistance, which can prevent a further increase in resistance by creating a ceiling effect. A possible interpretation is that alcohol acts to inhibit LNv neurons, which in turn increases resistance to alcohol and causes tolerance. This interpretation is supported by the fact that alcohol is a well-known potentiation of the inhibitory GABAreceptor (Lobo & Harris, 2008) and that GABAreceptors are known to modulate the release of PDF in *Drosophila* through the fly GABAreceptor homolog Rdl (Agosto *et al*., 2008; Parisky *et al*., 2008).

Interestingly, when we sought to silence the activity of LNv using different genetic tools to either electrically shunt action potentials (using the inward rectifier potassium channel Kir2.1) or inhibit synaptic fusion (using the tetanus toxin light chain peptide), we obtained two different results. When LNv neurons were electrically silenced using Kir2.1, flies maintained the ability to acquire tolerance. However, when we used TeTxLC to alter neurotransmitter release, flies lost the ability to acquire tolerance. While contradicting, these results are not surprising. These two methods of silencing neurons can impact distinct aspects of neural physiology and affect different behavioral processes. On the one hand, TeTxLC has been shown to directly interfere with synaptic transmission (Sweeney *et al*., 1995) and neuropeptide release (Ding *et al*., 2019). Silencing of LNv neurons using TeTxLC has been shown to have a modest effect on locomotor rhythmicity but severely disrupts morning anticipation in a manner that resembles the *Pdf* ^01^ mutation (Jaumouillé *et al*., 2021; Kaneko *et al*., 2000). On the other hand, the expression of Kir2.1 is known to reduce excitability by dampening electrical signals (Baines *et al*., 2001). Reducing membrane excitability through expression Kir2.1 within the LNv neurons results in severe behavioral arrhythmicity and loss of rhythmic molecular oscillations (Nitabach *et al*., 2002; Nitabach *et al*., 2005). To further explore the effect of the activity of LNv neurons on alcohol neuroadaptation, we also used over-expression of the sodium channel NaChBac to enhance the activity of PDF-expressing neurons. While this manipulation has been shown to affect rhythmicity significantly, it did not affect the ability of flies to acquire alcohol tolerance, suggesting again that the alcohol acts through the inhibition of PDF signaling rather than through the activation of PDF signaling. However, at this point, we do not know whether alcohol is acting directly on PDF release or through its effect on upstream or downstream signaling proteins.

Indeed, there are several potential mechanisms by which alcohol can affect PDF signaling. This includes the possibility that modulation of the alcohol response occurs through direct inhibition of either synaptic or somatic PDF release. Somatic PDF release has been shown to occur in response to chemical rather than electrical signals (Klose *et al*., 2021). This is consistent with the fact that inhibition of LNv neurons with TeTxLC was more effective at blocking alcohol tolerance than inhibition with Kir2.1. Another possibility is that alcohol is blocking downstream targets such as the PDF receptor (PDFR). This G protein-coupled receptor (GPCR) is specifically responsive to PDF and is expressed in a wide range of neurons, including a small subset of circadian pacemakers (Hyun *et al*., 2005; Lear *et al*., 2005; Mertens *et al*., 2005). Indeed, alterations in PDF signaling in the LNv neurons have been shown to occur through effects on PDF auto-receptors (Klose & Shaw, 2021). Moreover, alcohol exposure is known to affect the trafficking and function of many GPCRs that are associated with aberrant behavioral responses to alcohol (Luessen *et al*., 2019).

With these results in mind, we focused on uncovering the neuronal mechanisms of sleep by looking again at PDF-releasing LNv neurons as possible candidates for regulating this process. First, we established the effect alcohol has on sleep in wild-type flies. We found three critical aspects of sleep that were disturbed after alcohol exposure: (1) Alcohol exposure promotes daytime sleep. This effect is associated with a decrease in the number of sleep episodes and an increase in the mean sleep episode duration, suggesting that alcohol might play a role in consolidating sleep during the day. (2) Alcohol affects the structure of sleep during the night. Even though the total amount of sleep during the night is not affected by alcohol, we see clear differences in sleep during the first half of the night when compared to the second half of the night in alcohol-exposed flies. This effect is displayed in our analysis of the night sleep ratio. Alcohol primarily reduced sleep in the first half of the night but increased sleep in the second half. While it is unclear to us why this is happening, it suggests that the two halves correspond to distinct sleep phases and should be considered separately. The observed sleep pattern suggests that alcohol might reduce sleep quality during the first half of the night, promoting a homeostatic sleep rebound during the second half of the night and the daytime sleep phase. The phenomenon where organisms sleep more after a period of sleep deprivation to compensate for lost sleep is a well-established feature of sleep (Allada *et al*., 2017). (3) Alcohol disrupts morning anticipation. This effect is most likely connected with the increased sleep displayed by alcohol-treated flies during the last half of the night. The longer the flies sleep into the night, the lower the activity displayed in anticipation of the morning. Interestingly, lack of morning anticipation is one of the hallmark phenotypes of *Pdf ^01^* mutant flies, which again implicates the neuropeptide PDF in an alcohol-induced response (Grima *et al*., 2004; Renn *et al*., 1999; Stoleru *et al*., 2004). Close inspection of the role of PDF in alcohol-induced dysregulation confirmed that PDF is involved in the alcohol response. While we did not see a significant effect of the *Pdf ^01^* mutation on the alcohol-induced increase in total daytime sleep duration, we did see suppression of the alcohol effect on the number and duration of sleep episodes. This suggests that PDF is only partially responsible for the effects of alcohol on daytime sleep. In contrast, during the night, the *Pdf ^01^* mutation interfered with both the alcohol-induced disruption of sleep structure and morning anticipation, suggesting that the PDF plays a primary role in modulating sleep architecture during the night. One effect of alcohol that was not observed in wild-type flies but became evident in the *Pdf ^01^* mutant was the significant increase in total night-time sleep induced by alcohol. This effect was manifested only during the first night after exposure (Figure 2C) and stemmed mainly from the fact that the *Pdf ^01^* mutant had severely reduced sleep during the first night. The alcohol exposure seemed to rescue the deficit. One plausible explanation for this result is that alcohol suppresses the anxiety or stress associated with placing flies in a novel and perhaps even uncomfortable environment —the activity monitor tube.

In humans, sleep is often reduced when sleeping in unfamiliar surroundings in what is called the first night effect (Agnew *et al*., 1966). This effect is also evident in flies. Albeit subtle, it usually manifests as a gradual increase in the total sleep after flies are first placed in the activity monitors (Figure 2B). In *Pdf ^01^* mutants, however, this phenomenon is more severe during the first night (Figure 3B). While the exact role of PDF in this phenomenon has not been investigated, there is a well-studied connection between the PDF-producing LNv neurons and stress responses. One of the targets of the LNv neurons is the DN1 neuronal cluster. These neurons, while also part of the core clock cells in the *Drosophila* brain, have recently been associated with controlling stress and anxiety by releasing another peptide: the Diuretic Hormone 31 (DH31).

There is a significant amount of evidence that suggests that alcohol and sleep interact with each other through mechanisms related to stress. In humans, the hypothalamic-pituitary-adrenal axis and the extra-hypothalamic brain stress axis are under circadian control and regulate stress-related behavioral and neuroendocrine responses (Herman *et al*., 2016; Nader *et al*., 2010). These regions are also affected by alcohol (Stephens & Wand, 2012). From a molecular standpoint, evidence suggests that circadian genes such as CLOCK and CRY regulate the HPA axis and the secretion of hormones and neuropeptides associated with stress (Koch *et al*., 2017; Nader *et al*., 2010). Nevertheless, it is important to note that findings from Drosophila do not always extrapolate to mammalian systems and vice versa. While there is a strong conservation between the two at the molecular level, this conservation does not necessarily translate at the system level.

While there are still many unanswered questions regarding the role of the PDF neuropeptide in the relationship between alcohol and sleep, our results have shed light on potential mechanistic interactions. We propose that alcohol suppresses LNv signaling, resulting in a cascade of events that disrupt many aspects of sleep, potentially implicating stress-related mechanisms as well. Our research has also promoted the use of the *Drosophila* model to study the relationship between alcohol and sleep, which opens many new avenues of research related to this question.

## Author Contributions

M.E.R., J.L.A, and A.G. designed the study; M.E.R. and G.A. performed alcohol and sleep experiments; M.E.R., N.L.F, J.L.A., and A.G. analyzed the data; M.E.R., N.L.F, J.L.A., and A.G. wrote/reviewed the manuscript.

## Acknowledgments

We want to thank all members of the Ghezzi lab for their continuous input during the development of this study.

## References

Agnew HW, Webb WB, Williams RL (1966) The first night effect: an EEG study of sleep. Psychophysiology 2:263–266. PMID: 5903579.

Agosto J, Choi JC, Parisky KM, Stilwell G, Rosbash M, Griffith LC (2008) Modulation of GABAA receptor desensitization uncouples sleep onset and maintenance in Drosophila. Nat Neurosci 11:354–359. PMID: 18223647.

Allada R, Cirelli C, Sehgal A (2017) Molecular Mechanisms of Sleep Homeostasis in Flies and Mammals. Cold Spring Harb Perspect Biol 9 PMID: 28432135.

Artiushin G, Sehgal A (2017) The Drosophila circuitry of sleep-wake regulation. Curr Opin Neurobiol 44:243–250. PMID: 28366532.

Atkinson NS, Robertson GA, Ganetzky B (1991) A component of calcium-activated potassium channels encoded by the Drosophila slo locus. Science 253:551–555. PMID: 1857984.

Baines RA, Uhler JP, Thompson A, Sweeney ST, Bate M (2001) Altered electrical properties in Drosophila neurons developing without synaptic transmission. J Neurosci 21:1523–1531. PMID: 11222642.

Barber AF, Fong SY, Kolesnik A, Fetchko M, Sehgal A (2021) Drosophila clock cells use multiple mechanisms to transmit time-of-day signals in the brain. Proc Natl Acad Sci U S A 118 PMID: 33658368.

Brand AH, Manoukian AS, Perrimon N (1994) Ectopic expression in Drosophila. Methods Cell Biol 44:635–654. PMID: 7707973.

Brower KJ (2001) Alcohol’s effects on sleep in alcoholics. Alcohol Res Health 25:110–125. PMID: 11584550.

Brower KJ, Perron BE (2010) Prevalence and correlates of withdrawal-related insomnia among adults with alcohol dependence: results from a national survey. Am J Addict 19:238–244. PMID: 20525030.

Cabrero P, Radford JC, Broderick KE, Costes L, Veenstra JA, Spana EP, Davies SA, Dow JA (2002) The Dh gene of Drosophila melanogaster encodes a diuretic peptide that acts through cyclic AMP. J Exp Biol 205:3799–3807. PMID: 12432004.

Cannell E, Dornan AJ, Halberg KA, Terhzaz S, Dow JAT, Davies SA (2016) The corticotropin-releasing factor-like diuretic hormone 44 (DH44) and kinin neuropeptides modulate desiccation and starvation tolerance in Drosophila melanogaster. Peptides 80:96–107. PMID: 26896569.

Cavanaugh DJ, Geratowski JD, Wooltorton JR, Spaethling JM, Hector CE, Zheng X, Johnson EC, Eberwine JH, Sehgal A (2014) Identification of a circadian output circuit for rest:activity rhythms in Drosophila. Cell 157:689–701. PMID: 24766812.

Ceriani MF, Hogenesch JB, Yanovsky M, Panda S, Straume M, Kay SA (2002) Genome-wide expression analysis in Drosophila reveals genes controlling circadian behavior. J Neurosci 22:9305–9319.

Chung BY, Kilman VL, Keath JR, Pitman JL, Allada R (2009) The GABA(A) receptor RDL acts in peptidergic PDF neurons to promote sleep in Drosophila. Curr Biol 19:386–390. PMID: 19230663.

Cowmeadow RB, Krishnan HR, Ghezzi A, Al’Hasan YM, Wang YZ, Atkinson NS (2006) Ethanol tolerance caused by slowpoke induction in Drosophila. Alcohol Clin Exp Res 30:745–753. PMID: 16634842.

Daan S, Beersma DG, Borbély AA (1984) Timing of human sleep: recovery process gated by a circadian pacemaker. Am J Physiol 246:R161–83. PMID: 6696142.

Ding K, Han Y, Seid TW, Buser C, Karigo T, Zhang S, Dickman DK, Anderson DJ (2019) Imaging neuropeptide release at synapses with a genetically engineered reporter. Elife 8 PMID: 31241464.

Dube SR, Miller JW, Brown DW, Giles WH, Felitti VJ, Dong M, Anda RF (2006) Adverse childhood experiences and the association with ever using alcohol and initiating alcohol use during adolescence. J Adolesc Health 38:444.e1–10. PMID: 16549308.

Dubowy C, Sehgal A (2017) Circadian Rhythms and Sleep in Drosophila melanogaster. Genetics 205:1373–1397. PMID: 28360128.

Fernandez MP, Chu J, Villella A, Atkinson N, Kay SA, Ceriani MF (2007) Impaired clock output by altered connectivity in the circadian network. Proc Natl Acad Sci U S A 104:5650–5655. PMID: 17369364.

Furuya K, Milchak RJ, Schegg KM, Zhang J, Tobe SS, Coast GM, Schooley DA (2000) Cockroach diuretic hormones: characterization of a calcitonin-like peptide in insects. Proc Natl Acad Sci U S A 97:6469–6474. PMID: 10841553.

Geissmann Q, Garcia Rodriguez L, Beckwith EJ, Gilestro GF (2019) Rethomics: An R framework to analyse high-throughput behavioural data. PLoS One 14:e0209331. PMID: 30650089.

Ghezzi A, Atkinson NS (2011) Homeostatic control of neural activity: a Drosophila model for drug tolerance and dependence. Int Rev Neurobiol 99:23–50. PMID: 21906535.

Ghezzi A, Al-Hasan YM, Krishnan HR, Wang Y, Atkinson NS (2013) Functional mapping of the neuronal substrates for drug tolerance in Drosophila. Behav Genet 43:227–240. PMID: 23371357.

Ghezzi A, Krishnan HR, Atkinson NS (2014) Susceptibility to ethanol withdrawal seizures is produced by BK channel gene expression. Addict Biol 19:332–337. PMID: 22734584.

Goda T, Tang X, Umezaki Y, Chu ML, Kunst M, Nitabach MNN, Hamada FN (2016) Drosophila DH31 Neuropeptide and PDF Receptor Regulate Night-Onset Temperature Preference. J Neurosci 36:11739–11754. PMID: 27852781.

Grima B, Chélot E, Xia R, Rouyer F (2004) Morning and evening peaks of activity rely on different clock neurons of the Drosophila brain. Nature 431:869–873. PMID: 15483616.

Harrisingh MC, Wu Y, Lnenicka GA, Nitabach MN (2007) Intracellular Ca2+ regulates free-running circadian clock oscillation in vivo. J Neurosci 27:12489–12499. PMID: 18003827.

Hector CE, Bretz CA, Zhao Y, Johnson EC (2009) Functional differences between two CRF-related diuretic hormone receptors in Drosophila. J Exp Biol 212:3142–3147. PMID: 19749107.

Helfrich-Förster C (1995) The period clock gene is expressed in central nervous system neurons which also produce a neuropeptide that reveals the projections of circadian pacemaker cells within the brain of Drosophila melanogaster. Proc Natl Acad Sci U S A 92:612–616. PMID: 7831339.

Hendricks JC, Finn SM, Panckeri KA, Chavkin J, Williams JA, Sehgal A, Pack AI (2000) Rest in Drosophila is a sleep-like state. Neuron 25:129–138. PMID: 10707978.

Herman JP, McKlveen JM, Ghosal S, Kopp B, Wulsin A, Makinson R, Scheimann J, Myers B (2016) Regulation of the Hypothalamic-Pituitary-Adrenocortical Stress Response. Compr Physiol 6:603–621. PMID: 27065163.

Hodge JJ (2009) Ion channels to inactivate neurons in Drosophila. Front Mol Neurosci 2:13. PMID: 19750193.

Hyun S, Lee Y, Hong ST, Bang S, Paik D, Kang J, Shin J, Lee J, Jeon K, Hwang S, Bae E, Kim J (2005) Drosophila GPCR Han is a receptor for the circadian clock neuropeptide PDF. Neuron 48:267–278. PMID: 16242407.

Jaumouillé E, Koch R, Nagoshi E (2021) Uncovering the Roles of Clocks and Neural Transmission in the Resilience of Drosophila Circadian Network. Front Physiol 12:663339. PMID: 34122135.

Johns DC, Marx R, Mains RE, O’Rourke B, Marbán E (1999) Inducible genetic suppression of neuronal excitability. J Neurosci 19:1691–1697. PMID: 10024355.

Johnson EC, Shafer OT, Trigg JS, Park J, Schooley DA, Dow JA, Taghert PH (2005) A novel diuretic hormone receptor in Drosophila: evidence for conservation of CGRP signaling. J Exp Biol 208:1239–1246. PMID: 15781884.

Kaneko M, Park JH, Cheng Y, Hardin PE, Hall JC (2000) Disruption of synaptic transmission or clock-gene-product oscillations in circadian pacemaker cells of Drosophila cause abnormal behavioral rhythms. J Neurobiol 43:207–233. PMID: 10842235.

King AN, Sehgal A (2020) Molecular and circuit mechanisms mediating circadian clock output in the Drosophila brain. Eur J Neurosci 51:268–281. PMID: 30059181.

Klose MK, Bruchez MP, Deitcher DL, Levitan ES (2021) Temporally and spatially partitioned neuropeptide release from individual clock neurons. Proc Natl Acad Sci U S A 118 PMID: 33875606.

Klose MK, Shaw PJ (2021) Sleep drive reconfigures wake-promoting clock circuitry to regulate adaptive behavior. PLoS Biol 19:e3001324. PMID: 34191802.

Koch CE, Leinweber B, Drengberg BC, Blaum C, Oster H (2017) Interaction between circadian rhythms and stress. Neurobiol Stress 6:57–67. PMID: 28229109.

Koob GF, Le Moal M (1997) Drug abuse: hedonic homeostatic dysregulation. Science 278:52–58. PMID: 9311926.

Koob GF, Le Moal M (2001) Drug addiction, dysregulation of reward, and allostasis. Neuropsychopharmacology 24:97–129. PMID: 11120394.

Kunst M, Hughes ME, Raccuglia D, Felix M, Li M, Barnett G, Duah J, Nitabach MN (2014) Calcitonin gene-related peptide neurons mediate sleep-specific circadian output in Drosophila. Curr Biol 24:2652–2664. PMID: 25455031.

Lear BC, Merrill CE, Lin JM, Schroeder A, Zhang L, Allada R (2005) A G protein-coupled receptor, groom-of-PDF, is required for PDF neuron action in circadian behavior. Neuron 48:221–227. PMID: 16242403.

Littleton J (1998) Neurochemical mechanisms underlying alcohol withdrawal. Alcohol Health Res World 22:13–24. PMID: 15706728.

Lobo IA, Harris RA (2008) GABA(A) receptors and alcohol. Pharmacol Biochem Behav 90:90–94. PMID: 18423561.

Luan H, Lemon WC, Peabody NC, Pohl JB, Zelensky PK, Wang D, Nitabach MN, Holmes TC, White BH (2006) Functional dissection of a neuronal network required for cuticle tanning and wing expansion in Drosophila. J Neurosci 26:573–584. PMID: 16407556.

Luessen DJ, Sun H, McGinnis MM, Hagstrom M, Marrs G, McCool BA, Chen R (2019) Acute ethanol exposure reduces serotonin receptor 1A internalization by increasing ubiquitination and degradation of β-arrestin2. J Biol Chem 294:14068–14080. PMID: 31366729.

Menegazzi P, Beer K, Grebler V, Schlichting M, Schubert FK, Helfrich-Förster C (2020) A Functional Clock Within the Main Morning and Evening Neurons of D. melanogaster Is Not Sufficient for Wild-Type Locomotor Activity Under Changing Day Length. Front Physiol 11:229. PMID: 32273848.

Mertens I, Vandingenen A, Johnson EC, Shafer OT, Li W, Trigg JS, De Loof A, Schoofs L, Taghert PH (2005) PDF receptor signaling in Drosophila contributes to both circadian and geotactic behaviors. Neuron 48:213–219. PMID: 16242402.

Nader N, Chrousos GP, Kino T (2010) Interactions of the circadian CLOCK system and the HPA axis. Trends Endocrinol Metab 21:277–286. PMID: 20106676.

Nitabach MN, Blau J, Holmes TC (2002) Electrical silencing of Drosophila pacemaker neurons stops the free-running circadian clock. Cell 109:485–495. PMID: 12086605.

Nitabach MN, Sheeba V, Vera DA, Blau J, Holmes TC (2005) Membrane electrical excitability is necessary for the free-running larval Drosophila circadian clock. J Neurobiol 62:1–13. PMID: 15389695.

Okamoto-Mizuno K, Mizuno K (2012) Effects of thermal environment on sleep and circadian rhythm. J Physiol Anthropol 31:14. PMID: 22738673.

Parisky KM, Agosto J, Pulver SR, Shang Y, Kuklin E, Hodge JJ, Kang K, Kang K, Liu X, Garrity PA, Rosbash M, Griffith LC (2008) PDF cells are a GABA-responsive wake-promoting component of the Drosophila sleep circuit. Neuron 60:672–682. PMID: 19038223.

Park A, Ghezzi A, Wijesekera TP, Atkinson NS (2017) Genetics and genomics of alcohol responses in Drosophila. Neuropharmacology 122:22–35. PMID: 28161376.

Park JH, Hall JC (1998) Isolation and chronobiological analysis of a neuropeptide pigment-dispersing factor gene in Drosophila melanogaster. J Biol Rhythms 13:219–228. PMID: 9615286.

Pohl JB, Ghezzi A, Lew LK, Robles RB, Cormack L, Atkinson NS (2013) Circadian genes differentially affect tolerance to ethanol in Drosophila. Alcohol Clin Exp Res 37:1862–1871. PMID: 23808628.

Ren D, Navarro B, Xu H, Yue L, Shi Q, Clapham DE (2001) A prokaryotic voltage-gated sodium channel. Science 294:2372–2375. PMID: 11743207.

Renn SC, Park JH, Rosbash M, Hall JC, Taghert PH (1999) A pdf neuropeptide gene mutation and ablation of PDF neurons each cause severe abnormalities of behavioral circadian rhythms in Drosophila. Cell 99:791–802. PMID: 10619432.

Roehrs T, Roth T (1995) Alcohol-Induced Sleepiness and Memory Function. Alcohol Health Res World 19:130–135. PMID: 31798081.

Rothenfluh A, Troutwine BR, Ghezzi A, Atkinson NS. The genetics of alcohol responses of invertebrate model systems. In: Noronha ABC, Cui C, Harris RA, Crabbe JC, eds. Neurobiology of alcohol dependence. Amsterdam, Netherlands: Elsevier; 2014: 463–491.

SAMHSA (2017) 2017 National Survey of Drug Use and Health (NSDUH) Releases. https://www.samhsagov/data/release/2017-national-survey-drug-use-and-health-nsduh-releases

Scholz H, Ramond J, Singh CM, Heberlein U (2000) Functional ethanol tolerance in Drosophila. Neuron 28:261–271. PMID: 11086999.

Shafer OT, Yao Z (2014) Pigment-Dispersing Factor Signaling and Circadian Rhythms in Insect Locomotor Activity. Curr Opin Insect Sci 1:73–80. PMID: 25386391.

Shang Y, Griffith LC, Rosbash M (2008) Light-arousal and circadian photoreception circuits intersect at the large PDF cells of the Drosophila brain. Proc Natl Acad Sci U S A 105:19587–19594. PMID: 19060186.

Shaw PJ, Cirelli C, Greenspan RJ, Tononi G (2000) Correlates of sleep and waking in Drosophila melanogaster. Science 287:1834–1837. PMID: 10710313.

Sheeba V, Sharma VK, Gu H, Chou YT, O’Dowd DK, Holmes TC (2008a) Pigment dispersing factor-dependent and -independent circadian locomotor behavioral rhythms. J Neurosci 28:217–227. PMID: 18171939.

Sheeba V, Fogle KJ, Kaneko M, Rashid S, Chou YT, Sharma VK, Holmes TC (2008b) Large ventral lateral neurons modulate arousal and sleep in Drosophila. Curr Biol 18:1537–1545. PMID: 18771923.

Sink KS, Walker DL, Yang Y, Davis M (2011) Calcitonin gene-related peptide in the bed nucleus of the stria terminalis produces an anxiety-like pattern of behavior and increases neural activation in anxiety-related structures. J Neurosci 31:1802–1810. PMID: 21289190.

Stephens MA, Wand G (2012) Stress and the HPA axis: role of glucocorticoids in alcohol dependence. Alcohol Res 34:468–483. PMID: 23584113.

Stoleru D, Peng Y, Agosto J, Rosbash M (2004) Coupled oscillators control morning and evening locomotor behaviour of Drosophila. Nature 431:862–868. PMID: 15483615.

Sweeney ST, Broadie K, Keane J, Niemann H, O’Kane CJ (1995) Targeted expression of tetanus toxin light chain in Drosophila specifically eliminates synaptic transmission and causes behavioral defects. Neuron 14:341–351. PMID: 7857643.

Thakkar MM, Sharma R, Sahota P (2015) Alcohol disrupts sleep homeostasis. Alcohol 49:299–310. PMID: 25499829.

Todd WD, Venner A, Anaclet C, Broadhurst RY, De Luca R, Bandaru SS, Issokson L, Hablitz LM, Cravetchi O, Arrigoni E, Campbell JN, Allen CN, Olson DP, Fuller PM (2020) Suprachiasmatic VIP neurons are required for normal circadian rhythmicity and comprised of molecularly distinct subpopulations. Nat Commun 11:4410. PMID: 32879310.

White BH, Osterwalder TP, Yoon KS, Joiner WJ, Whim MD, Kaczmarek LK, Keshishian H (2001) Targeted attenuation of electrical activity in Drosophila using a genetically modified K(+) channel. Neuron 31:699–711. PMID: 11567611.

